# Internal desynchrony of the circadian clock system in middle-aged mice under social jet lag-like conditions

**DOI:** 10.1101/2025.10.01.679730

**Authors:** Saki Fukushima, Dan Yang, Tatsumi Morita, Kaito Kurogi, Ryohei Matsuo, Tiantian Ma, Keisuke Ikegami, Mitsuhiro Furuse, Shinobu Yasuo

## Abstract

Social jet lag (SJL) refers to the discrepancy in sleep patterns between weekdays and weekends, leading to a misalignment between the internal clock and social time. In this study, we investigated the effects of weekly shifts in light-dark (LD) conditions: two days per week with 6-h delayed LD cycles (simulating Saturday and Sunday), followed by a 6-h advance on Monday. Core body temperature rhythms rapidly entrained to the delayed LD cycles on weekends, and these delayed rhythms persisted even after the LD cycle was advanced on Monday. In contrast, plasma corticosterone rhythms on Mondays were aligned with the LD cycle but exhibited reduced amplitude. In the livers of SJL mice on Monday, the expression rhythms of *Per1*, *Per2*, and *Hsp70* were delayed by 3-5 h compared to that in the controls, whereas *Rev-erbα* expression rhythms remained comparable to those of the controls. The expression of lipid and glucose metabolism-related genes in the liver showed either delayed rhythms or no significant changes. To determine whether the dissociation of gene expression rhythms resulted from gene-specific responses to circadian body temperature and hormonal signals, we conducted ex vivo culture experiments using mouse liver slices. High-temperature stimulation induced *Per2* and *Hsp70* expression, while dexamethasone induced *Per1* expression. High temperature and dexamethasone affected distinct sets of metabolic genes, whereas insulin induced only minor changes. Moreover, these responses were strongly influenced by the age and light exposure of the mice. We also examined the effect of weekly housing by providing environmental enrichment (EE), which had minimal impact on circadian parameters but promoted anti-aging effects on bone density and behavior. Overall, our findings indicate that weekly shifts in LD cycles induce internal desynchronization within the hepatic molecular clock and metabolic pathways by uncoupling core body temperature rhythms, hormonal rhythms, and gene-specific responses to stimuli.

## Introduction

The circadian clock regulates approximately 24-h rhythms of behavioral and physiological processes. In mammals, the master clock is located in the suprachiasmatic nucleus (SCN), which coordinates circadian oscillations in both the central nervous system and peripheral organs. One of the key mediators of peripheral clock resetting is core body temperature, whose circadian rhythms are regulated by the SCN^1,2^. The entrainment of peripheral clocks to body temperature involves heat shock factor 1 (HSF1) and HSF1-driven heat shock proteins (HSP)^1,3,4^. Rhythms of hepatic HSPs are observed even in the absence of a functional clock^4^, indicating that they are primarily regulated by systemic body temperature rhythms, independently of the local clock. Circulating glucocorticoids also contribute to the resetting of peripheral clocks. These hormones are rhythmically secreted from the adrenal gland in an SCN-dependent manner via the sympathetic nervous system and help synchronize peripheral clocks^2^. At the molecular level, circadian rhythms are generated by transcriptional-translational feedback loops composed of clock genes such as *Per1*, *Per2*, *Bmal1, Clock,* and *Reb-erbα*^5^. The molecular clock is tightly coupled to physiological functions through the transcriptional regulation of clock-controlled genes. For instance, the cell cycle is linked to the circadian clock, as CLOCK and BMAL1 directly regulate the expression of cell cycle genes *c-Myc* and *Wee1*^6,7^. Peripheral clocks generate rhythmicity in glucose and lipid metabolism^8,9^, while hippocampal clocks play critical roles in neurogenesis, synaptic plasticity, and cognition^10,11^.

Aberrant lighting conditions, such as exposure to bright light at night or frequent shifts in light–dark (LD) cycles, lead to circadian desynchronization or disruption. Chronic circadian desynchronization is widely recognized to be closely associated with an increased risk of diseases, including obesity, diabetes, psychiatric disorders, and neurological diseases^12,13^. In animal studies, circadian disruption has been induced by exposure to constant light (LL), chronic jet lag (CJL), or forced desynchronization protocols^14^. These treatments cause clock desynchronization within cells and organs; consequently, clock-controlled physiological functions become misaligned or are lost altogether. These effects may be exacerbated by aging, as the circadian clock in aged mice exhibits a blunted response to shifts in LD cycles^15,16^. Repeated advances in LD cycles every several days have been reported to increase mortality in mice and to accelerate immune senescence^17,18^.

Social jet lag (SJL) refers to a discrepancy in sleep patterns between weekdays and weekends in humans, resulting in a misalignment between the internal circadian clock and social time^19,20^. SJL has been associated with increased body mass index and a higher risk of prediabetes and type 2 diabetes^19,21,22^. In animal studies, the weekly pattern of circadian delay and advance observed in SJL was replicated by repeated shifts of LD cycles over a seven-day schedule: a 6-h delay for two days (i.e., experimental Saturday and Sunday), followed by a return to the standard cycle with 6-h advances over five days (i.e., experimental Monday to Friday)^23,24^. In SJL model mice, activity rhythms and Per2 protein rhythms in both the central and peripheral clocks were delayed throughout the week, accompanied by impaired cognitive function^23^. Considering that the re-entrainment speeds of peripheral clocks to acute LD advances are highly tissue- and gene-specific^25,26^, the SJL-like model may exhibit clock desynchronization among molecular clocks, metabolic rhythms, and systemic rhythm mediators. However, few studies have demonstrated the dissociation of expression rhythms in clock- and metabolism-associated genes within a single organ or the dissociation of systemic cues such as core body temperature and hormones.

This study aimed to investigate how SJL-like conditions involving weekly shifts in the LD cycle induce desynchronization of the circadian clock system, as well as alterations in behavioral and metabolic parameters. We used middle-aged mice, as aging is associated with slower re-entrainment of shifted circadian rhythms to environmental cycles,^16^ and is expected to exacerbate circadian desynchronization. Circadian clock disruption leads to cognitive and metabolic impairments, primarily related to dysfunction in the hippocampus and liver^8,15,27^. To compare the sensitivity of the hippocampus and liver to abnormal LD conditions, we first examined the effects of two well-established circadian disturbance paradigms—LL and CJL—on gene expression in these tissues and on behavioral outcomes prior to the SJL-like experiments. In the SJL experiment, we evaluated the effects of weekly changes in housing conditions by providing environmental enrichment (EE) for two days per week. Chronic EE is known to enhance cognitive ability, reduce stress responses, and mitigate disease pathologies^28^. Weekly EE mimics the variation in comfort between weekdays and weekends, based on the hypothesis that stress levels may interact with weekly shifts in the circadian clock.

## Results

### SJL-like conditions impact body composition, grading, behavior, and hepatic clock

We first compared the effect of LL and CJL on aging-related behaviors. The CJL group showed increased exploratory activity in the open field test (OFT) (*p* = 0.02), whereas the LL group exhibited reduced distance traveled in the center zone (*p* = 0.0171; Supplementary Fig. 1A–D). No significant effects of LL or CJL were detected in the rotarod test and object recognition test (ORT) (Supplementary Fig. 1E–J), the tests for evaluation of motor coordination and cognition. We also examined the effects of LL and CJL on the temporal expression of clock genes in the liver and hippocampus. Hsp70 was included as a marker of systemic, body temperature–dependent regulation. In the liver, the rhythms of clock genes (*Per1, Per2, Baml1*, and *Rev-erbα)* and *Hsp70* were significantly altered by both LL and CJL exposure (Supplementary Fig. 2A). In contrast to the varied changes observed in the liver, the hippocampus showed only minor alterations in the expression of clock genes and *Hsp70* under LL or CJL conditions (Supplementary Fig. 2B). Thus, we decided to analyze affective behaviors, metabolic/somatic outcomes, and hepatic gene expression rhythms in the following SJL-like experiments.

For SJL-like conditions using LD cycle shifts, the lights-on and lights-off times were advanced by 6 h on day 1, delayed by 6 h on day 6, and returned to the original cycle by 6 h on day 1 of the following cycle (weekly LD treatment, LD; Fig. 1A). These patterns induce phase delays and advances in the circadian clock^23^, thereby mimicking working days (days 1– 5, Monday–Friday) and weekends (days 6 and 7, Saturday and Sunday) in people. For weekly EE, mice were transferred to EE cages during the light phase on day 6 and returned to standard cages on day 1 of the subsequent cycle. Additional group was exposed to weekly LD and EE housing (LD-EE). Both LD and EE treatments increased body weight (*p* < 0.0001) over experimental periods without significant interaction between them (Fig. 1B). At the final point, LD treatment increased body weight (*p* = 0.0008) and epididymal fat weight (*p* = 0.0108), independent of weekly EE (Fig. 1B–D). Trabecular and total bone densities—but not cortical bone density—increased at week 16 with weekly EE (*p* = 0.0463 and 0.0156, respectively), regardless of lighting conditions (Fig. 1E–G). The total grading score during week 4–16 was significantly reduced in EE mice (*p* = 0.0221) but not in LD and LD-EE mice (Supplementary Fig. 3). Hair loss inhibition was observed only in EE mice for each grading item (*p* = 0.0029) (Supplementary Fig. 3). These findings suggest that EE treatment has an anti-aging effect, which is partially attenuated by weekly shifts in the LD cycle.

**Figure 1.**
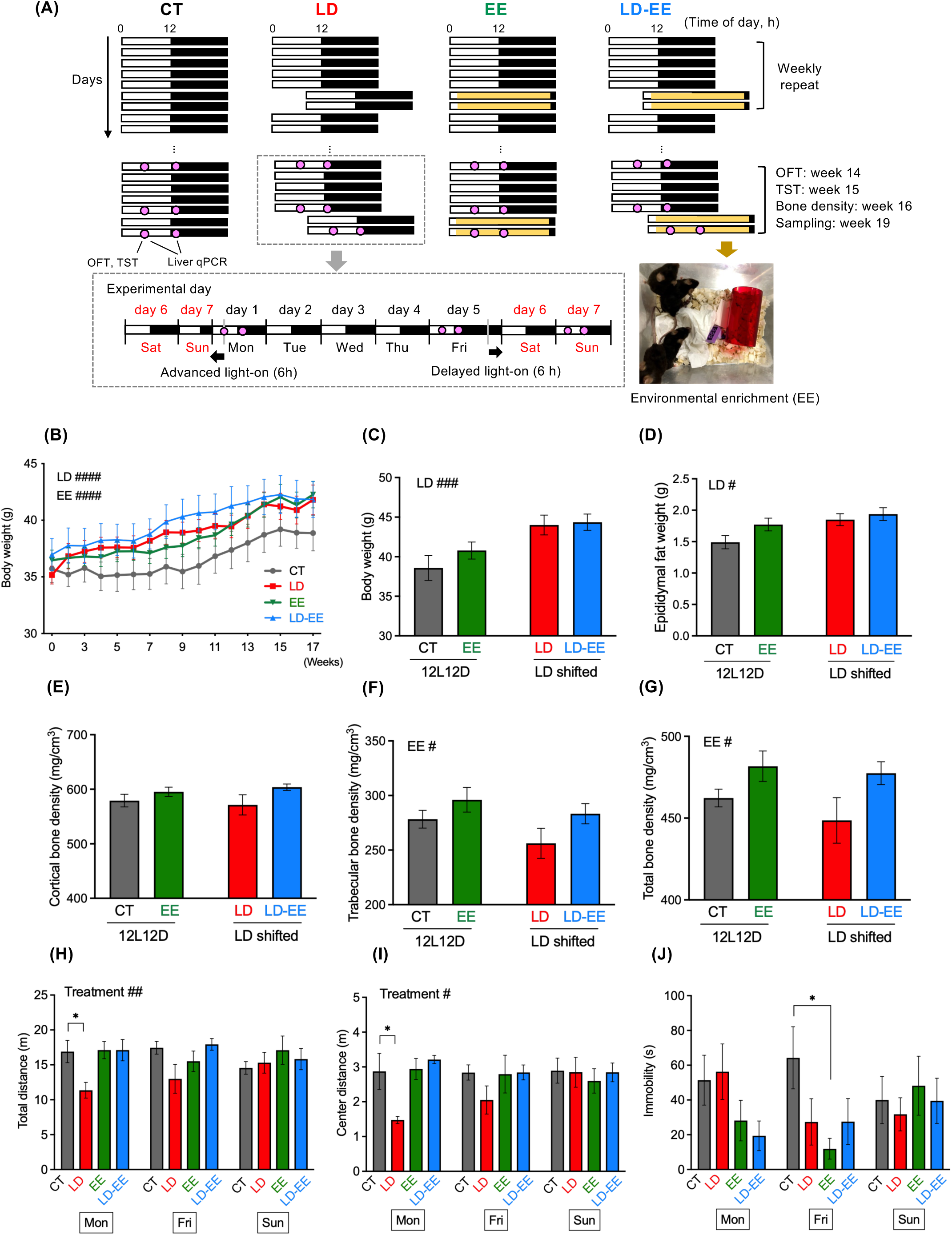
SJL-like conditions affect body weight, bone density, and anxiety- and depression-like behavior. (A) Experimental timelines for control (CT), weekly shifts of light-dark cycles (LD), weekly housing in environmental enrichment (EE), and a combination of LD and EE (LD-EE). For the LD and LD-EE groups, the light-dark cycle was 6 h delayed on simulated Friday (Day 5) by prolonging the dark phase, followed by a 6 h advance on simulated Monday by shortening the dark phase. For the EE and LD-EE groups, mice were housed in EE cages for two days (simulated weekends) in a weekly cycle. Open field test (OFT), tail suspension test (TST), bone density evaluation by computerized tomography, and liver tissue sampling were conducted at indicated week. (B–D) Effect of SJL-like conditions on temporal changes in body weight (B), final body weight (C), and epididymal fat weight (D). (E–G) Effect of SJL-like conditions on cortical (E), trabecular (F), and total bone density (G) in the femurs (n = 9). (H– J) Effect of SJL-like conditions on behavior in OFT (H and I) and TST (J) (n = 8–9 per group). Data are shown as mean ± standard error of the mean (S.E.M). #*p* < 0.05, ##*p* < 0.01, ###*p* < 0.001, two-way ANOVA; \**p* < 0.05, Bonferroni’s multiple comparison test.

In the OFT, the total distance traveled and the distance traveled in the center were significantly lower with LD mice on Mondays (*p* = 0.0277 and 0.0256, respectively) (Fig. 1H and I). This effect was not observed in LD-EE mice, suggesting a preventive effect of weekly EE on LD shifts–induced increase in anxiety-like behavior. Immobility in the TST was significantly suppressed in EE mice on Fridays (*p* = 0.0248) (Fig. 1J). This effect was not pronounced in LD-EE mice, indicating that LD shifts abolished the antidepressant effect of weekly EE.

Next, we examined the expression of clock genes (*Per1* and *Rev-erbα*), the clock-controlled cell cycle regulatory gene (*Wee1*), and *Hsp70* in the liver at two time points per day on Mondays, Fridays, and Sundays. Diurnal variations in *Per1*, *Rev-erbα*, and *Wee1* expression were stable among the four treatment groups (control (CT), LD, EE, and LD-EE) on Mondays and Fridays (Fig. 2). In contrast, the diurnal expression patterns of these genes were markedly altered on Sundays in the LD and LD-EE groups (Fig. 2). For *Hsp70*, the diurnal expression pattern was stable across the four treatment groups on Fridays and Sundays, but altered patterns were observed on Mondays in the LD and LD-EE groups (Fig. 2). These results suggest that the expression rhythms of the hepatic clock and clock-control genes are not entrained to LD cycle delays on Saturdays and Sundays, whereas *Hsp70* expression rhythms are entrained to the cycle but fail to entrain to LD advances on Mondays.

**Figure 2.**
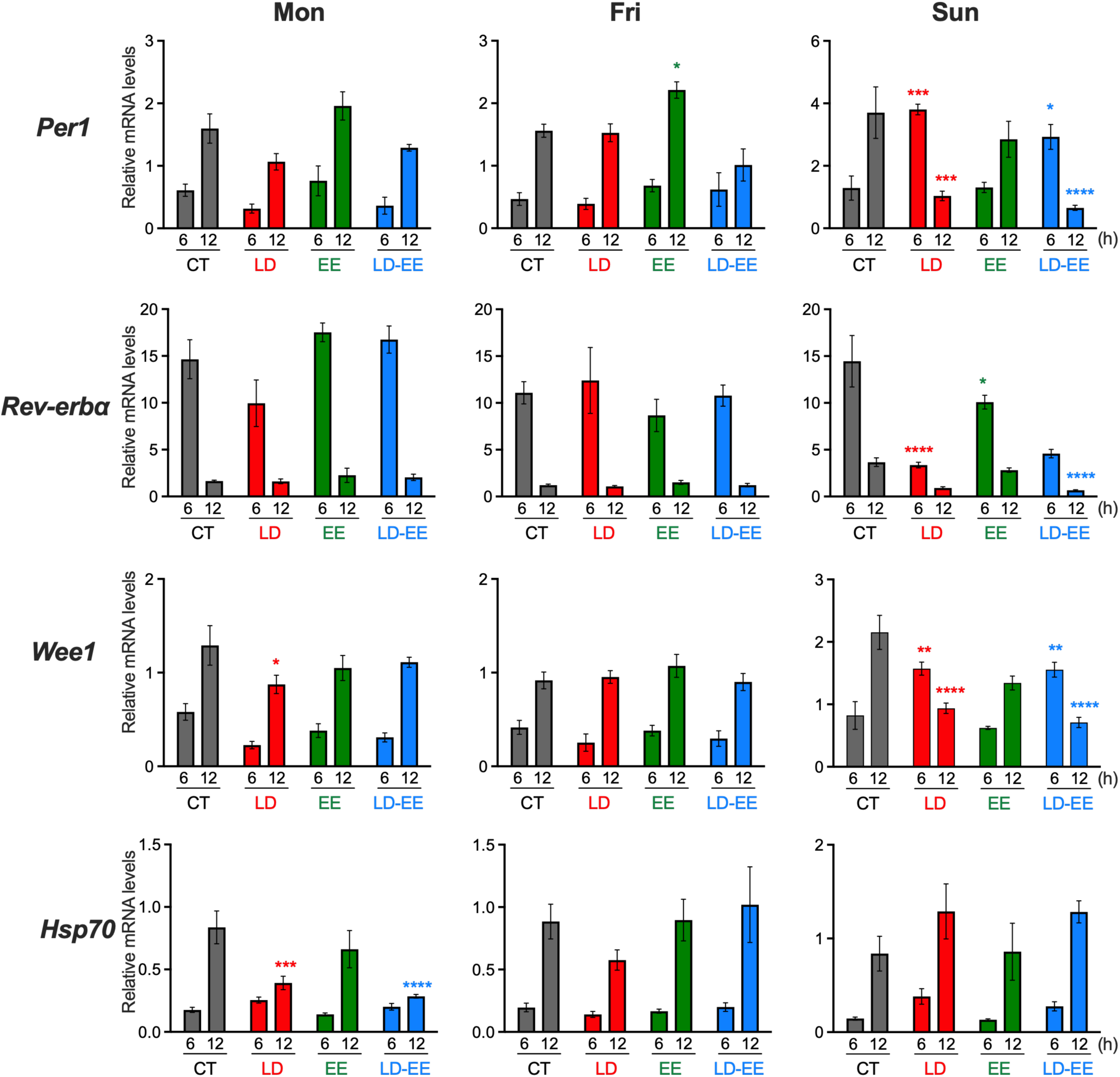
SJL-like conditions alter day-night variations in liver gene expression in a gene-specific manner. Expression at two time points is shown for control (CT), weekly shifts of light-dark cycles (LD), weekly housing in environmental enrichment (EE), and a combination of LD and EE (LD-EE) on simulated Monday, Friday, and Sunday (n = 3–4). Data are shown as mean ± standard error of the mean (S.E.M). \**p* < 0.05, \*\**p* < 0.01, \*\*\**p* < 0.001, \*\*\*\**p* < 0.0001, Bonferroni’s multiple comparison test, comparison with values at respective time point in CT group.

### SJL-like conditions with weekly LD shifts induce dissociation between plasma corticosterone and body temperature rhythms

To confirm the gene-specific expression responses to SJL-like conditions with weekly LD shifts, we analyzed six time points on a simulated Monday (Fig. 3A). To examine the diurnal patterns of systemic circadian regulators, plasma corticosterone levels and core body temperature rhythms were analyzed. Body weight significantly increased in LD mice (*p* = 0.0062) (Fig. 3B), whereas food intake was not significantly altered (CT: 3.08 ± 0.19 g; LD: 3.226 ± 0.15 g). Plasma corticosterone levels in CT group showed significant diurnal variations, with a peak at 10:00 (*p* = 0.0039), which were blunted in the LD group with non-significant variations (Fig. 3C). Heatmaps and temporal plots of core body temperature showed stable rhythmicity throughout the week in CT mice (Fig. 3D and E). In contrast, LD mice exhibited weekly delays in daily peak body temperature, particularly on Mondays and Tuesdays (Fig. 3F and G). The average body temperature peaks across the three weeks remained stable in CT mice, whereas daily variation was evident in LD mice (Fig. 3H and I). In LD mice, body temperature peaks on Mondays (*p* = 0.0012), Tuesdays (*p* = 0.0056), and Sundays (*p* < 0.0001) were significantly delayed compared to those on Fridays. Body temperature amplitude also fluctuated within a week in LD mice, being significantly reduced on Mondays (*p* = 0.0012) and Tuesdays (*p* = 0.0052) compared with Sundays (Fig. 3J). These findings indicate that plasma corticosterone and body temperature rhythms became dissociated on Mondays under a weekly LD-shift schedule.

**Figure 3.**
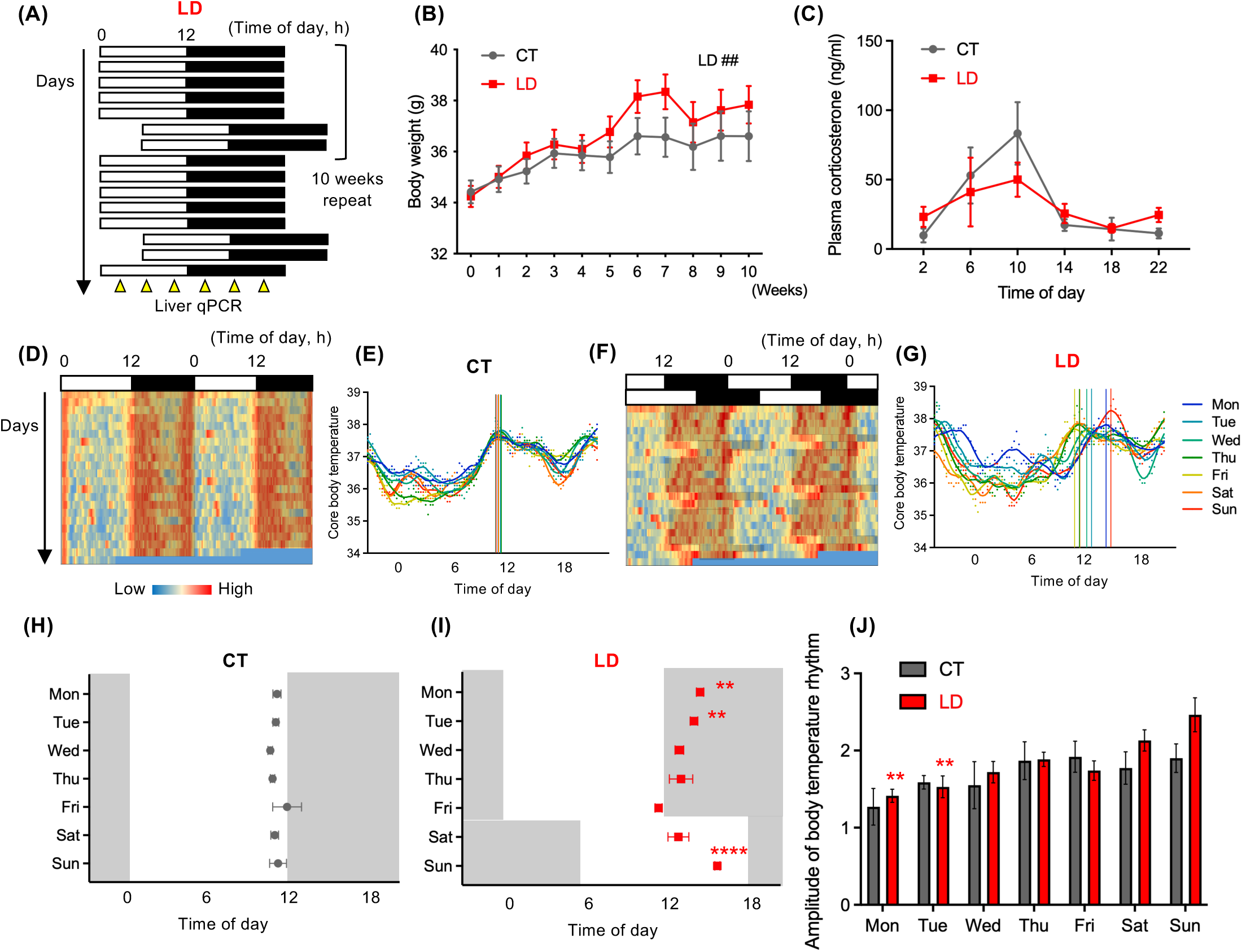
SJL-like conditions with weekly shifts in light-dark cycles (LD) induce desynchronization between plasma corticosterone rhythms and core body temperature rhythms. (A) Experimental timeline for LD treatment. (B) Temporal changes in body weight in the control (CT) and LD groups (n = 24). ##*p* < 0.01, two-way ANOVA. (C) Plasma corticosterone rhythms on simulated Mondays under LD treatment (n = 4). (D–G) Heatmaps and daily variations of core body temperature in the CT (D, E) and LD (F, G) groups. Data for representative animals are shown for each group. Peaks for each day are indicated by colored lines. (H, I) Peak timing of core body temperature rhythms in CT (H) and LD (I) groups (n = 4). \*\**p* < 0.01, \*\*\*\**p* < 0.0001, Dunnett’s multiple comparison test, comparison with values on simulated Friday. (J) Amplitude of core body temperature rhythms each day (n = 4). \*\**p* < 0.01, Bonferroni’s multiple comparison test, comparison with values on simulated Sunday. Data are shown as mean ± standard error of the mean (S.E.M).

### SJL-like conditions with weekly LD shifts induce internal desynchronization in the hepatic clock and metabolism

We examined the expression rhythms of clock and clock-controlled genes related to the cell cycle, lipid metabolism, and glucose metabolism, alongside *Hsp70*, in the liver on Monday following weekly LD shifts mimicking SJL (Fig. 3A). Among the clock genes examined, acrophases of *Per1* and *Per2* expression were delayed by 4.637 and 4.136 h in LD mice, whereas delays of *Bmal1* and *Rev-erbα* expression were limited compared to them (Fig. 4A, Supplementary Table 1). *Hsp70* and *Wee1* expression exhibited 3.729 and 2.796 h delay, whereas other clock-controlled cell cycle–associated genes (*p53* and *c-Myc*) were largely unaffected (Fig. 4B, Supplementary Table 1). Similarly, several clock-controlled metabolic genes, including *Cyp7a1* showed delayed expression rhythms, whereas other genes showed little to no change (Fig. 4D). To analyze the protein-metabolizing conditions, we analyzed diurnal variations in plasma free amino acid levels. In CT mice, 24 amino acids were clustered into three major groups (Fig. 4E). Diurnal variations in amino acids in each cluster were visualized using normalized data (Fig. 4E). In LD mice, the peaks of clusters 2 were delayed by approximately 4 h compared with those in CT mice. In contrast, the amino acids in cluster 3 retained patterns similar to those in CT mice. These findings suggest that hepatic gene expression and metabolic processes undergo internal desynchronization on Mondays under SJL-like conditions, characterized by a dissociation between delayed and retained components.

**Figure 4.**
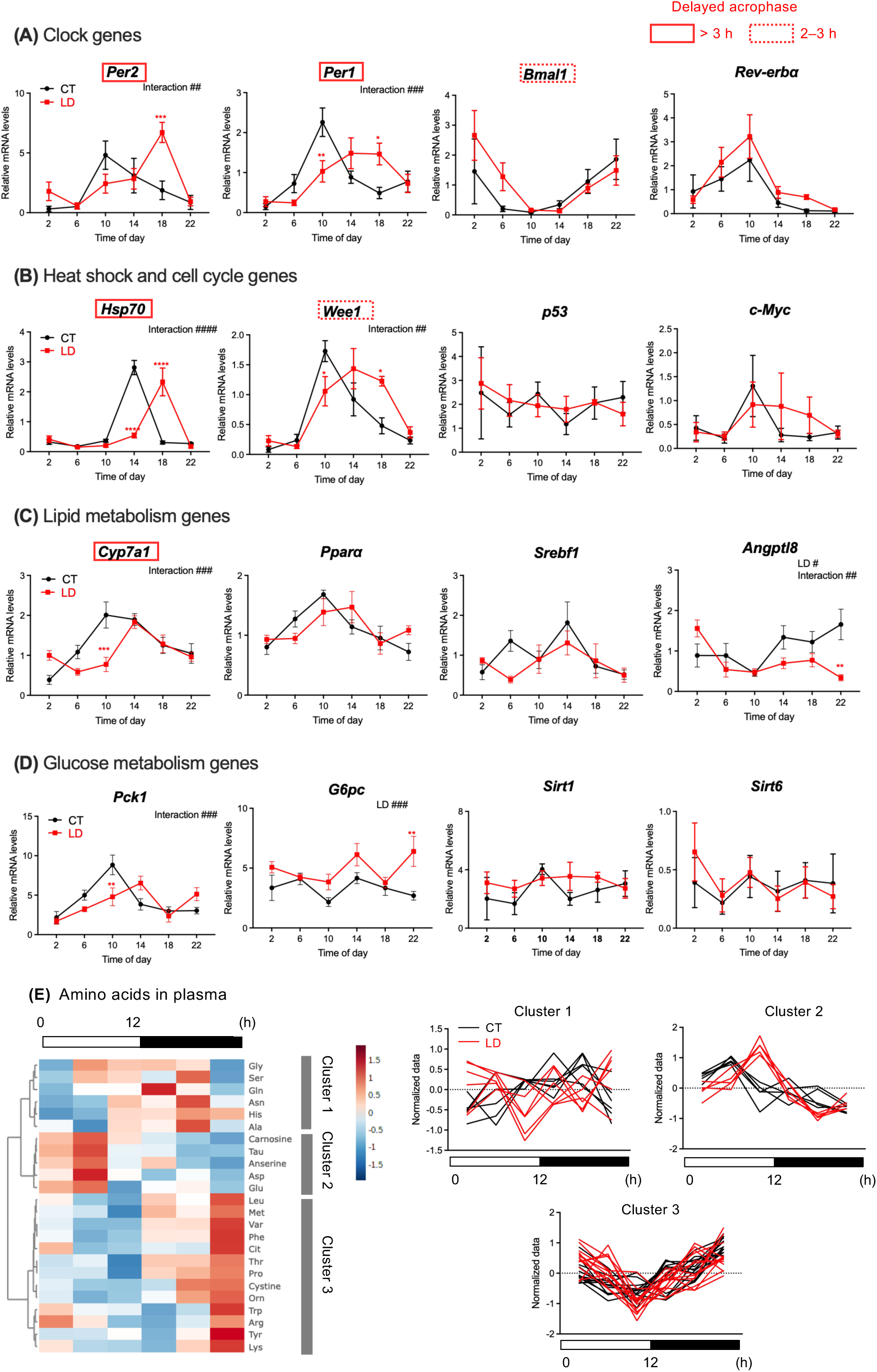
SJL-like conditions with weekly shifts in light-dark cycles (LD) induce desynchronization of hepatic gene expression rhythms. (A–D) Temporal expression of clock genes (A), heat shock and cell cycle genes (B), lipid metabolism genes (C), and glucose metabolism genes (D) on simulated Monday in the control (CT) and LD groups (n = 3–4). Red squares in gene names indicate genes with delayed peaks in the LD group, based on cosinor analyses (delayed acrophases more than 3 h: solid lines; 2–3 h: dashed lines). (E) Heatmap showing temporal changes in plasma free amino acid levels in CT groups. Based on normalized data in each cluster, along with corresponding data in LD groups, temporal changes in amino acids are shown in right panels. Data are shown as mean ± standard error of the mean (S.E.M). \**p* < 0.05, \*\**p* < 0.01, \*\*\**p* < 0.001, \*\*\*\**p* < 0.0001, Bonferroni’s multiple comparison test.

### High temperature, dexamethasone, and insulin induce gene-specific responses in liver slices

The experiments described above revealed dissociation between systemic signals—such as body temperature and hormonal rhythms—and the expression rhythms of hepatic clock and metabolic genes. Based on these findings, we hypothesized that weekly shifts in the LD cycle induce internal desynchronization within the hepatic molecular clock and metabolic pathways through the misalignment of core body temperature rhythms, hormonal rhythms, and gene-specific responses to these stimuli. To test this hypothesis, we performed ex vivo cultures using liver slices from middle-aged mice to analyze gene-specific responses following 3- and 6-h treatment with high temperature, dexamethasone, a synthetic glucocorticoid, and insulin (Fig. 5A). High temperature induced the expression of *Per2* (6 h), *Hsp70* (3 and 6 h), and *Pck1* (6 h) (Fig. 5B), all of which exhibited delayed peak rhythms in LD mice on Mondays (Fig. 4A, B, D). Dexamethasone (6 h) induced the expression of *Per1*, *Pparα*, and *G6pc* (Fig. 5B), which showed either blunted rhythms or insignificant alterations in LD mice (Fig. 4A). In contrast, insulin induced minimal changes in the expression of all examined genes (Fig. 5B). Taken together, high temperature and dexamethasone affect distinct gene sets, suggesting that the dissociation of gene expression rhythms on Mondays in LD mice arises from gene-specific responses to systemic signals.

**Figure 5.**
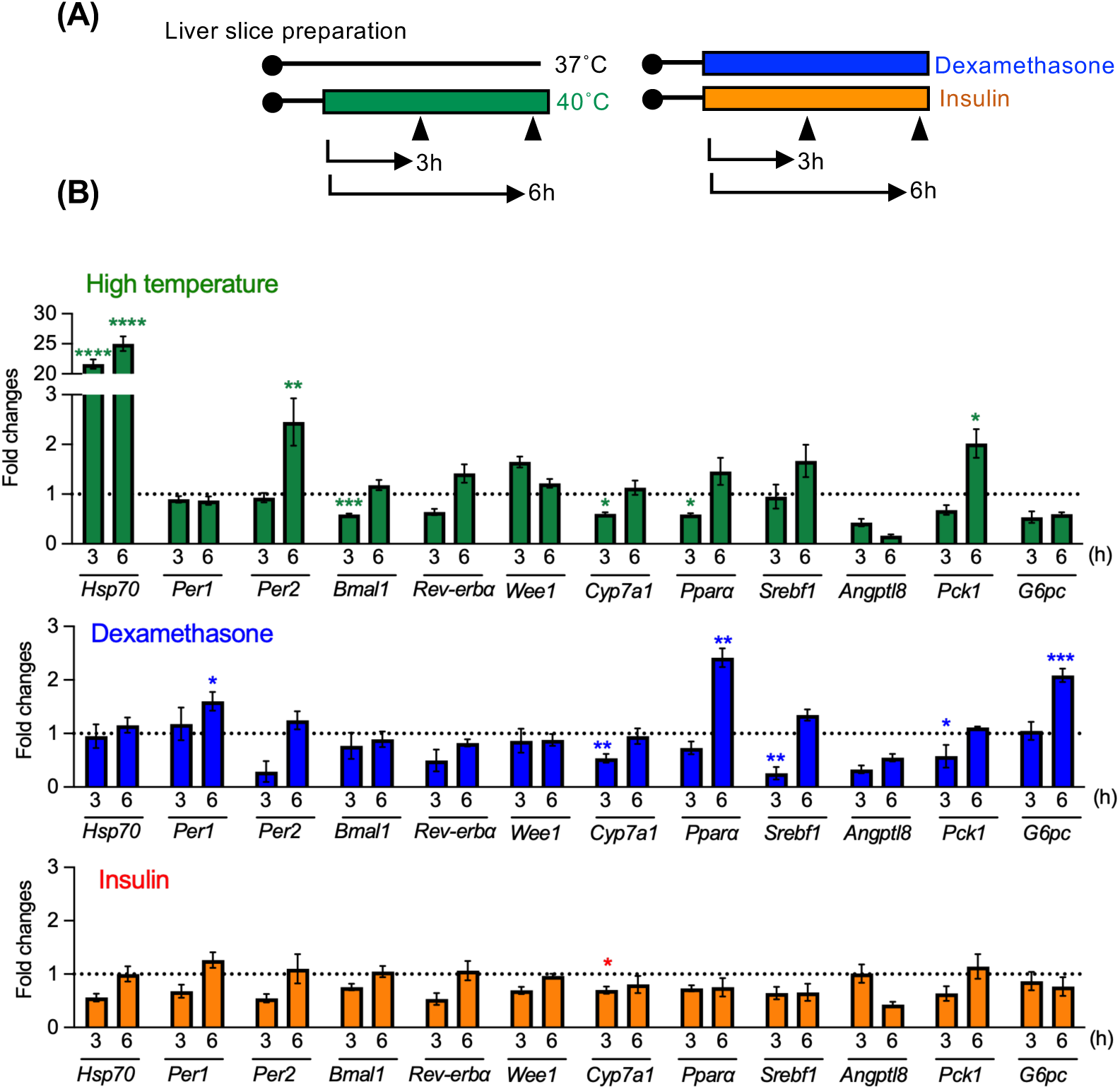
Gene expression in liver slices responds to high temperature, dexamethasone, and insulin stimulation in a gene-specific manner. (A) Experimental timeline of slice preparation, stimulation, and tissue collection. Liver slices were prepared from middle-aged mice maintained under a light/dark (LD) cycle of 12:12. (B) Effect of high temperature, dexamethasone, and insulin on the expression of heat-sensitive *Hsp70*, circadian clock genes, clock-controlled genes, and metabolic genes in liver slices (three replicates per treatment). Data are shown as fold changes relative to the expression levels in non-stimulated slices. Data are shown as mean ± standard error of the mean (S.E.M). \**p* < 0.05, \*\**p* < 0.01, \*\*\**p* < 0.001, \*\*\*\**p* < 0.0001, Bonferroni’s multiple comparison test, comparison with values in non-stimulated slices.

We further investigated whether aging and weekly shifts in the LD cycle affect gene expression. Liver slices were prepared from young and middle-aged mice maintained under 12L:12D (CT) or subjected to weekly shifts of the LD cycle and then treated with high temperature, dexamethasone, and insulin. Aging enhanced the responses of several clock and metabolic genes (*Per1*, *Per2*, *Cyp7a1*, and *Pck1*) to 3 h of insulin treatment in LD mice, whereas it blunted the induction of *Hsp70* expression by insulin (Fig. 6). Aging had minor or attenuated effects on gene responses to high temperature and dexamethasone (Fig. 6 and Supplementary Fig. 4 for 6 h stimulation). LD treatment markedly reduced the induction of *Hsp70* expression by high temperature in both young and middle-aged mice (Fig. 6 and Supplementary Fig. 4). The suppressive effect of LD exposure was also observed in response to 3 h of insulin treatment (Fig. 6) but not after 3 h of dexamethasone. Among the metabolic genes examined, *G6pc* was the most responsive to 6 h of dexamethasone and insulin treatment (Supplementary Fig. 4). Collectively, these findings suggest that gene-specific responses to systemic signals such as high temperature, dexamethasone, and insulin are strongly influenced by both aging and weekly shifts in the LD cycle.

**Figure 6.**
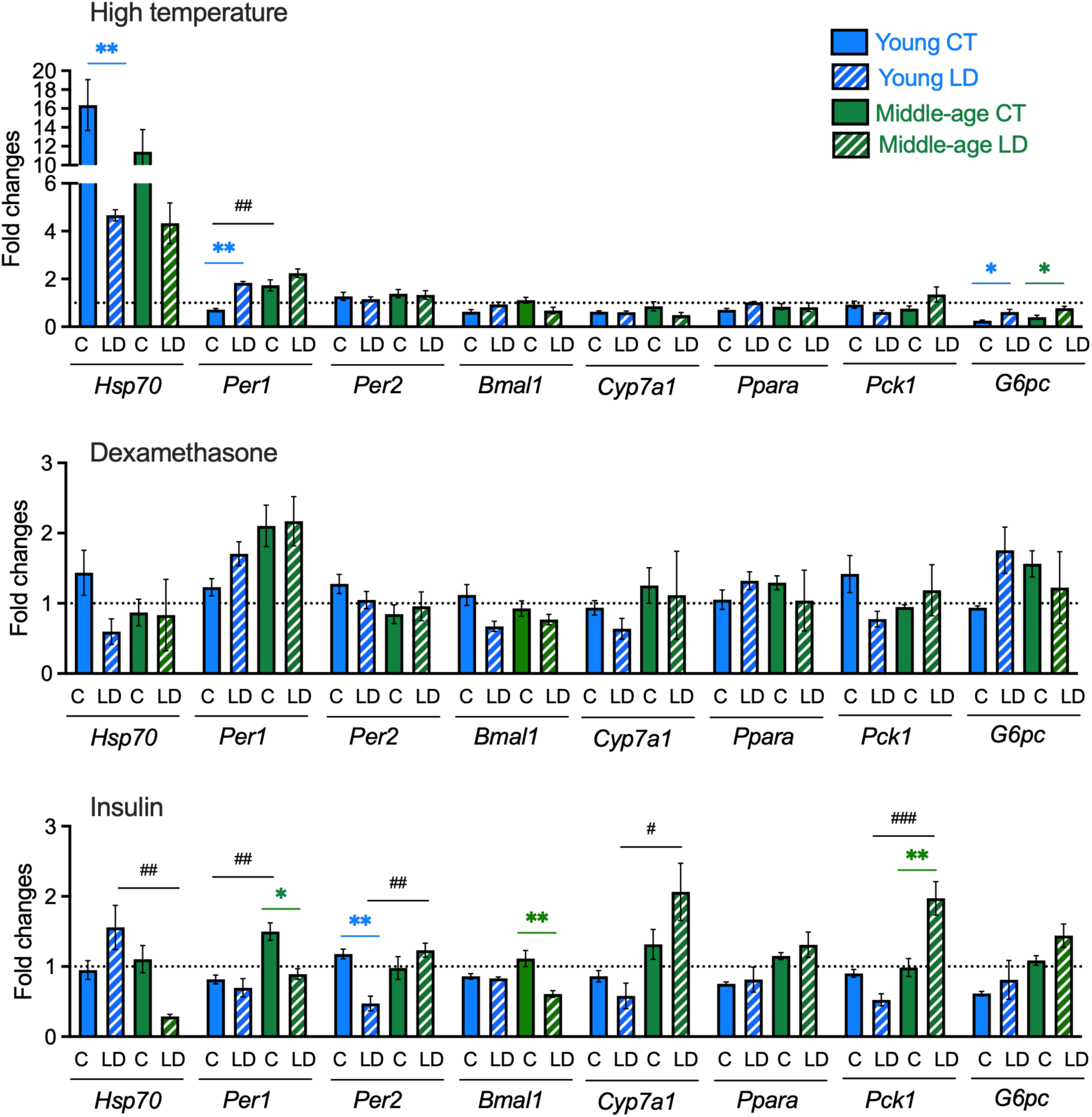
The responsiveness of gene expression in liver slices is altered by aging and lighting conditions of mice prior to preparation. Liver slices were prepared from young and middle-aged mice kept under LD 12:12 (CT) or weekly shifts of light-dark cycles (LD). Slices were stimulated with high temperature, dexamethasone, and insulin for 3 h and then analyzed for the expression of heat-sensitive *Hsp70*, circadian clock, and metabolic genes (three replicates per treatment). Data are shown as fold changes relative to the expression levels in non-stimulated slices. Data are shown as mean ± standard error of the mean (S.E.M). \**p* < 0.05, \*\**p* < 0.01, Bonferroni’s multiple comparison test, comparison between young CT and young LD and between middle-age CT and middle-age LD. #*p* < 0.05, ##*p* < 0.01, ###*p* < 0.001, Bonferroni’s multiple comparison test, comparison between the young and middle-aged CT groups and between the young and middle-aged LD groups.

## Discussion

In this study, we demonstrated that SJL-like conditions, involving weekly shifts in the LD cycle, induced desynchronization between core body temperature and glucocorticoid rhythms in middle-aged male mice. On simulated Mondays, core body temperature rhythms were delayed, resembling the delay observed in locomotor activity rhythms^23^, whereas glucocorticoid rhythms remained aligned with the LD cycle. These findings led us to hypothesize that systemic desynchronization promotes dissociation between hepatic clock components and their downstream pathways. We further identified dissociation in the expression rhythms of clock-, cell cycle-, and lipid/glucose metabolism–associated genes. One group of genes exhibited delayed rhythms, whereas another group remained mostly synchronized with the LD cycle. A similar dissociation was observed in plasma free amino acid rhythms, which reflect protein metabolism. Furthermore, liver slice culture experiments demonstrated gene-specific responses to either elevated temperatures or hormonal stimuli. These responses were significantly influenced by both age and prior lighting conditions. Taken together, weekly shifts in the LD cycle cause internal desynchronization within the liver molecular clock and clock-controlled pathways through the misalignment of systemic signals (Fig. 7). Indeed, the re-entrainment speed of peripheral clocks to abrupt LD cycle advances is highly tissue- and gene-specific^25^ and is closely associated with core body temperature rhythms^26^.

**Figure 7.**
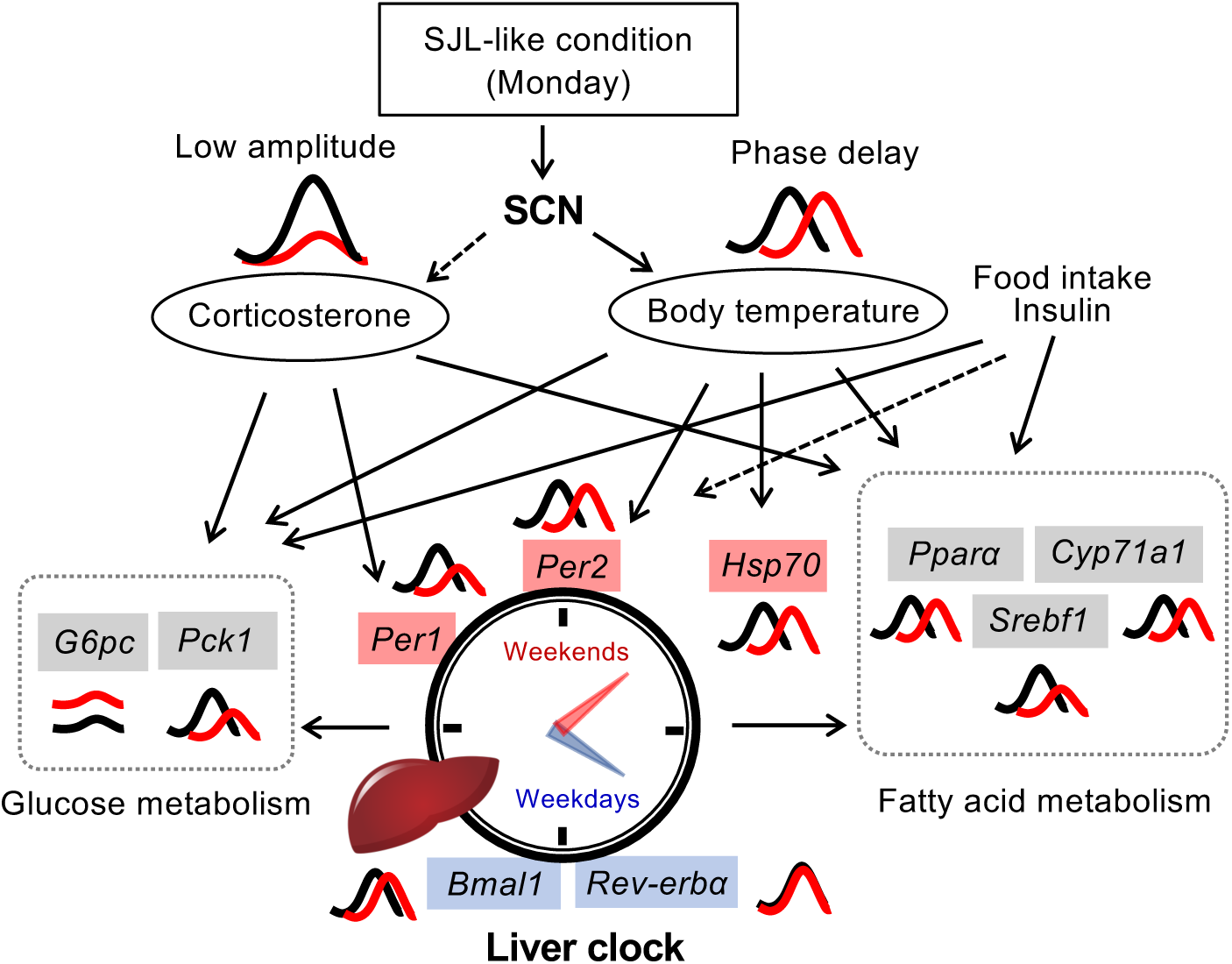
Schematic of the internal desynchronization proposed in the present study. Under SJL-like conditions, a 6 h delay of the light-dark cycle during two days each week (simulated weekends) followed by a 6 h advance of the cycle (simulated Mondays) results in a delay of the core body temperature rhythm, while the plasma corticosterone rhythm remains aligned (control: black waves; SJL-like condition: red waves). Food intake and insulin rhythms may also be altered. The dissociation of these systemic circadian signals affects the liver clock and associated metabolic pathways in a gene-specific manner. Molecular loops involving *Bmal1* and *Rev-erbα* maintain local clocks. Consequently, internal desynchronization occurs within the circadian clock system.

Under SJL-like conditions, the dissociation of day–night variation in clock gene expression in the liver was evident on simulated Sundays. The expression patterns of *Per1*, *Rev-erbα*, and *Wee1* on Sundays differed from those observed in control mice, whereas the pattern of *Hsp70* expression remained comparable to that of controls. Considering the rapid entrainment of core body temperature on Sundays in our study, the *Hsp70* expression pattern likely reflects full entrainment. In contrast, subtle or partial entrainment of *Per1* and *Rev-erbα* expression rhythms may have resulted in their expression being detected at misaligned time points. This slower entrainment is supported by the stable *Rev-erbα* expression rhythm observed on Mondays and its insensitivity to various stimuli in liver slice culture. A similar stability was observed in *Bmal1* expression, suggesting that the regulatory loop involving *Rev-erbα* and *Bmal1* contributes to the maintenance of hepatic clock function under systemic perturbations. However, the expression rhythm of *Per1* on Mondays was delayed and exhibited reduced amplitude. This finding is consistent with the high sensitivity of *Per1* to glucocorticoids and other signals, as demonstrated in this study. In mice exposed to repeated shifts in the LD cycle, the plasma corticosterone rhythm was dampened^29^, and resynchronization to an 8-h advanced LD cycle required seven days^30^. Therefore, corticosterone rhythms on Sundays were unlikely to be synchronized, and the rhythm peak likely persisted into Monday with reduced amplitude, as reflected in the observed *Per1* expression patterns.

Our study demonstrated largely delayed *Per1* and *Per2* expression rhythms in the liver on Mondays under SJL-like conditions, along with delayed expression of *Hsp70*. High-temperature stimulation induced *Per2* and *Hsp70* expression in liver slices, suggesting that body temperature regulates the oscillations in their expression. Previous studies have shown that circadian oscillation of hepatic Per2 production is modulated by both the molecular clock and HSF1-mediated temperature signaling^1,4,31^. Notably, *Per2* and *Hsp70* expression rhythms persist even in the absence of functional clock circuitry^4^. *Per2* expression in fibroblasts responds to heat-shock pulses in an HSF1-dependent manner^32^. However, a recent study reported that physiological temperature stimulation enhances Per2 translation without affecting *Per2* transcription^33^. This discrepancy may be explained by the age-dependent increase in *Per2* sensitivity observed in this study. We also demonstrated that aging amplifies the transcriptional response of *Per1* to high temperature and insulin stimulation. Aging reshapes the expression patterns of hepatic clock genes^34^, accompanied by changes in circadian clock oscillation, locomotor activity, and hormonal rhythms^35^. Age-dependent sensitization of clock genes to systemic signals may therefore contribute to the functional characteristics of the aged circadian clock.

Among the metabolic genes analyzed, the peak expression rhythms of *Cyp7a1* were significantly delayed on Mondays. Expression rhythms of *Pparα*, *and Pck1* also showed delayed tendency. Their expression is regulated by both the molecular clock^36–38^ and systemic signals, as evidenced by their gene-specific responses to elevated temperature and hormonal stimulation in this study. Other genes exhibited distinct regulatory patterns, such as the overall upregulation *of G6pc* or the minimal effect on *Srebf1* expression. Collectively, these findings suggest that internal desynchronization occurs within lipid and glucose metabolic pathways, which may in turn contribute to increased body weight. As the responses of several clock and metabolic genes to insulin stimulation were amplified by aging and SJL-like conditions, disrupted feeding rhythms are likely to play a central role in this desynchronization. Weekly shifts in food intake have been shown to induce phase shifts in peripheral clocks^23^. Further investigation is required to analyze food intake patterns and plasma insulin rhythms under SJL-like conditions.

Our analysis primarily focused on liver tissue, as hepatic clock rhythms were markedly dampened or phase-shifted under LL or CJL, whereas these treatments had minimal impact on the hippocampal clock, which is critical for memory and cognitive processes. Moreover, such light manipulations exerted minimal effect on behavioral tests strongly associated with aging, including the ORT and rotarod^39,40^. In contrast, studies in young mice have reported that LL and CJL impair the hippocampal clock, neuroplasticity, memory, and cognition^41,42^. This age-dependent discrepancy may be attributable to age-related dampening of hippocampal clock activity. For instance, a proteomic study showed that only 1.6% of proteins exhibited circadian rhythmicity in middle-aged mice (44–52 weeks old), compared with 15% in young mice (9–10 weeks old)^43^. The influence of aging on the hippocampal clock thus differs from that on the hepatic clock, where aged mice maintain oscillatory amplitudes comparable to those of young mice under both LD 12:12 and simulated jet lag conditions^44^. Given that aging is a major factor attenuating hippocampal rhythmicity and impairing cognitive function^45^, the adverse impact of aberrant lighting conditions is likely to be less pronounced in older animals.

We found that weekly exposure to EE increased bone density and decreased grading scores, suggesting an anti-aging effect of this intermittent EE paradigm. Previous studies have reported that continuous semi-natural or EE housing improves bone density and reduces alopecia in aged mice^46,47^, as well as consistently shows positive effects on aging-associated behaviors, including cognition and anxiety^48^. Notably, we observed that weekly shifts in the LD cycle increased anxiety-like behavior, consistent with previous findings^15,49^, and that this effect was counteracted by concurrent weekly EE housing. Conversely, depression-like behavior was reduced by weekly EE exposure, but this effect was abolished by LD cycle shifts. Together, these findings suggest that circadian misalignment and EE housing interact. Although the underlying neurobehavioral mechanisms remain unclear, our findings suggest that enhancing housing conditions through the introduction of environmental novelty and complexity on weekends may mitigate the negative behavioral consequences of SJL-induced circadian misalignment.

In conclusion, weekly shifts in the LD cycle lead to internal desynchronization of the hepatic molecular clock and associated metabolic pathways, driven by altered core body temperature rhythms, hormonal fluctuations, and gene-specific responses to these systemic cues. The extent of this desynchronization is strongly influenced by both age and light exposure. Moreover, weekly alterations in housing conditions affect age-related physiological parameters and behavioral outcomes. Collectively, these findings highlight the need for a comprehensive understanding of health disturbances under jet lag–like conditions, accounting for both biological rhythms and environmental factors that fluctuate weekly.

## Methods

### Animals

Male C57BL/6J mice were purchased from Japan SLC (Shizuoka, Japan) and housed in plastic cages in groups of three or four. Water and a standard laboratory rodent diet (MF; Oriental Yeast, Tokyo, Japan) were provided *ad libitum* and replenished at least once per week. Mice were maintained in a light-tight box placed in a temperature-controlled room (24 ± 1°C) under 12 h light (70 lx):12 h dark (12L:12D) conditions for at least two weeks prior to experimentation. The study was conducted in accordance with the Guidelines for Animal Experiments of the Faculty of Agriculture at Kyushu University, Law No. 105, and Notification No. 6 of the Japanese Government. All experimental procedures were approved by the Animal Care and Use Committee of Kyushu University (A19-212-0, A19-365-0, and A20-236-0).

### LL and CJL experiments

Experiments involving LL and CJL were conducted separately, using mice aged more than 30 weeks at the start of treatment. The control group was maintained under 12L:12D conditions, the LL group was exposed to continuous light conditions, and the CJL group underwent repeated 6-h advances of the LD cycle every two days, as previously described^50^. All groups were housed under standard conditions with bedding material. Behavioral testing was initiated at week 4 of the experiments. The OFT, ORT, and rotarod test were conducted sequentially, with intervals of several days between tests. After all tests were conducted, the control and CJL groups were transferred to constant darkness (DD) for two days and then euthanized using isoflurane gas at 0:00, 6:00, 12:00, or 18:00 based on circadian time (0:00 was defined as the light onset on the previous day; n = 3–5 per time point). In the control and CJL groups, the dark phase was initiated at 12:00 prior to the transition to DD. Mice in the LL group were euthanized at the same time points under LL conditions. Hippocampal and liver samples were harvested and stored in RNAlater (Thermo Fisher Scientific, Waltham, MA, USA) for quantitative PCR (qPCR) analysis.

### SJL-like experiments

Twenty-seven-week-old mice were divided into four treatment groups: control (CTL, 12L:12D), weekly shifts in the LD cycle, weekly exposure to environmental enrichment (EE), and a combination of LD and EE treatments (LD-EE) (Fig. 1A). To habituate the mice to the housing conditions, the EE and LD-EE groups were housed in EE cages equipped with a tunnel, two sheets of cardboard, nesting materials, and two block toys. The CTL and LD groups were housed under standard conditions with bedding material, under a 12L:12D cycle.

At 35 weeks of age, weekly cycles of LD conditions and EE housing were initiated. The LD and LD-EE groups were exposed to a weekly cycle (days 1–7) of light–dark shifts: on day 1 lights-on and lights-off times were advanced by 6 h by shortening the dark phase; this new cycle was maintained from days 2 to 5. On day 6, lights-on and lights-off times were delayed by 6 h by extending the dark phase, and this continued on day 7. The EE and LD-EE groups experienced repeated cycles of standard housing for five days (days 1–5), followed by two days (days 6 and 7) of EE housing. The CTL group remained under standard housing with 12L:12D conditions throughout the experiment, with gentle handling performed twice per week.

Grading scores^51^ were recorded at weeks 4, 8, and 16, and femoral bone density was measured using the Latheta LCT-100 system (Hitachi, Tokyo, Japan) at week 16. The OFT and TST were performed at weeks 14 and 15, respectively. Behavioral tests were conducted on days 1, 5, and 7 using separate cohorts of animals. Mice were euthanized at week 19 using isoflurane gas at 6:00 and 12:00 (n = 3–5 per time point), and liver samples were harvested and stored for qPCR analysis.

Another cohort of 35-week-old mice was divided into CTL and LD groups, with the LD group exposed to weekly shifts in the light–dark cycle, as described above. Six weeks after the start of the experiment, a temperature logger (diameter: 17 mm, height: 6 mm; KN Laboratories, Osaka, Japan) was implanted into the abdominal cavity under isoflurane anesthesia administered using an inhalation device (Natsume Seisakusho Co., Ltd., Tokyo, Japan). Following surgery, cephalexin (15 mg/kg) was dissolved in saline and injected subcutaneously. Core body temperature was recorded every 15 min until the end of the experiment. In week 10, mice were euthanized using isoflurane gas at 0:00, 6:00, 12:00, or 18:00 (n = 3–5 per time point). After decapitation, trunk blood was collected into heparinized tubes and centrifuged at 3000 × *g* for 10 min at 4°C to obtain plasma samples for corticosterone and amino acid analyses. Liver samples were harvested and stored for qPCR analysis.

### Liver ex vivo cultures

In the first experiment, mice over 9 months of age maintained under 12L:12D were used. The second experiment included both young (8-week-old) and middle-aged (over 9 months old) mice, which had been maintained under either 12L:12D or subjected to weekly shifts in the LD cycle, as described above, for more than 10 weeks. Mice were euthanized using isoflurane gas, and their livers were immersed in Hank’s balanced salt solution on ice at middle of light phases. Liver tissue was sliced into 250 µm sections using a tissue chopper (Cavey Laboratory Engineering Co Ltd, Surrey, UK) The slices were preincubated on a culture membrane (Millicell-CM PICM0RG50, Merck Millipore, Darmstadt, Germany) in a 35-mm petri dish and cultured in serum free Dulbecco’s modified Eagle’s medium (FUJIFILM Wako Pure Chemical Corporation, Osaka, Japan) supplemented with 1% penicillin / streptomycin in a 5 % CO_2_ incubator at 36°C for 1 h, then stimulated with either high temperature (39 °C), 100 nM dexamethasone (Merck Millipore, Burlington, MA, USA), a synthetic glucocorticoid, or 100 nM insulin (FUJIFILM Wako Pure Chemical Corporation). The slices were collected for qPCR analysis after 3 and 6 h of stimulation.

### Behavioral tests

The OFT was conducted in a white square box (40 × 40 cm) with 40 cm-high walls under 40 lx illumination. Each mouse was placed at the center of the apparatus and recorded for 5 min using a digital video camera. Open-field behavior was analyzed using the ANY-maze software (Stoelting Co., IL, USA), with the field divided into 25 squares (a 5 × 5 grid). The total distance moved, distance moved within the nine central grids, and time spent in the central grids (in 30-s bins) were automatically recorded by the software.

The ORT was performed the day after the OFT using the same box. During the familiarization session, each animal was allowed to move freely in the box containing a pair of identical objects (either centrifuge tubes or volumetric flasks) for 10 min. In the test session, conducted 20 min after, one of the familiar objects was replaced with a novel object, and the animal was left in the box for 5 min. The total time spent exploring each object was determined manually by the experimenter. The recognition index for novel object exploration was calculated as (N − F)/(N + F) × 100, where *F* is the time spent exploring the familiar object and *N* is the time spent exploring the novel object.

The accelerating rotarod test was conducted using a six-lane rotarod apparatus (MK-610A; Muromachi, Tokyo, Japan). The rotation speed was increased from 4 rpm to 40 rpm over a 5-min period. The latency to fall from the rotating rod was recorded, with a 6-min cutoff, across three trials at 1-h intervals over four consecutive days.

### Plasma corticosterone assay

Total plasma corticosterone levels were analyzed in duplicate using a corticosterone EIA kit (Cayman Chemicals, Ann Arbor, MI) according to the manufacturer’s protocol, with the exception of using 2.5% Steroid Displacement Reagent (Enzo Life Sciences, Farmingdale, NY, USA) during the plasma dilution step. The detection limit of the assay was 30 pg/ml. The intra- and interassay coefficients of variation were 2.8% and 8.9%, respectively.

### qPCR

Total RNA was extracted using RNAiso Plus (Takara, Shiga, Japan), according to the manufacturer’s protocol. cDNA was synthesized from 1 µg of total RNA using the PrimeScript RT Reagent Kit with gDNA Eraser (Takara). qPCR was performed on a Stratagene Mx3000P system (Agilent Technologies; Santa Clara, CA) with an initial denaturation step at 95°C for 30 s, followed by 40 cycles of amplification at 95°C for 5 s and 60°C for 30 s. Primer sequences are listed in Supplemental Table 2. mRNA levels were quantified based on threshold cycle values and compared with a standard curve. The calculated levels were normalized to the mRNA levels of *Hprt1* and *Rplp0*. A melting curve analysis was performed for each gene to confirm the specificity of the PCR conditions.

### Amino acid analyses

Amino acid levels were measured using fluorescence-based ultra-performance liquid chromatography. Tissue samples were homogenized in 0.2 M ice-cold perchloric acid containing 0.01 mM EDTA×2Na and kept on ice for 30 min to allow deproteinization. The samples were then centrifuged at 20,000 × *g* for 15 min at 0°C. Supernatants were filtered through a 0.2-µm filter (Millipore, Billerica, MA, USA) and adjusted to pH 3 with 1 M NaOH. The derivatized samples were analyzed using an ACQUITY UPLC system, consisting of a Waters Binary Solvent Manager, Sample Manager, FLR Detector, ACCQTAG^TM^ ULTRA C18 column (1.7 μm, 2.1 × 100 mm), and Empower 3 software (Waters, Milford, MA). The excitation and emission wavelengths for fluorescence detection were 350 nm and 450 nm, respectively.

### Statistical analysis

Body weight and data from the OFT and rotarod tests were analyzed using repeated measures two-way ANOVA, with treatment and weeks, days, or time as factors. Other data were analyzed using *t*-tests, one-way, or two-way ANOVA. Post-hoc analyses were performed using Bonferroni’s multiple comparisons test. For core body temperature, fluctuations each day across three weeks were averaged and smoothed (GraphPad Prism, San Diego, CA, USA). The peak time and amplitude of the smoothed rhythms were then compared among days or groups.

For evaluation of hepatic gene expression rhythms, cosinor analysis was performed based on linear harmonic regression using CircWave software (Roelof Hut, University of Groningen, version 1.4) with 0.05 for an assumed period of 24 h. This analysis allowed us to estimate the significance of circadian variations, acrophases as centers of gravity, and the amplitude from the peak to the middle of the fitted cosinor wave.

Hierarchical clustering and heatmap visualization of amino acid rhythms were conducted using MetaboAnalyst (http://www.metaboanalyst.ca). Data were normalized by median signal intensity, and auto-scaling was used to adjust scale differences.

For ex vivo data, fold changes in expression levels were calculated relative to unstimulated control tissues. In the first experiment, expression values for each stimulation were compared to control values using *t-*tests, while in the second experiment, fold changes across the four groups (young-12L:12D, young-LD, old-12L:12D, and old-LD) were compared using two-way ANOVA followed by Bonferroni’s multiple comparisons test. For all statistics, differences were considered statistically significant at *p* < 0.05.

## Acknowledgements

We would like to thank Editage (www.editage.com) for English language editing. This work was supported by the AMED under Grant Number JP24gm2010005h0001 and JST FOREST program, Japan (grant number: JPMJFR201D) to SY.

## Additional Information

We declare no conflicts of interest.

