## Supplementary Tables and Figures for "Internal desynchrony of the circadian clock system in middle-aged mice under social jet lag-like conditions"

**Supplementary Table 1** Evaluation of circadian rhythmicity of genes expression in the liver by cosinor analysis.

| | CT | | | LD | | | Acrophase delay |
|---|---|---|---|---|---|---|---|
| Gene | Acrophase | Amplitude | *p*-value | Acrophase | Amplitude | *p*-value | |
| *Per1* | 10.826 | 0.673 | < 0.001 | 15.464 | 0.709 | < 0.001 | 4.637 |
| *Per2* | 12.507 | 1.964 | 0.005 | 16.643 | 2.109 | 0.005 | 4.136 |
| *Bmal1* | 22.232 | 0.931 | 0.004 | 1.099 | 1.162 | < 0.001 | 2.867 |
| *Rev-erbα* | 7.906 | 0.997 | 0.005 | 9.049 | 1.353 | < 0.001 | 1.143 |
| *Hsp70* | 14.041 | 0.867 | < 0.001 | 17.770 | 0.745 | < 0.001 | 3.729 |
| *Wee1* | 11.837 | 0.674 | < 0.001 | 14.634 | 0.708 | < 0.001 | 2.796 |
| *p53* | 0.873 | | n.s. | 3.435 | | n.s. | |
| *c-Myc* | 9.561 | | n.s. | 13.452 | | n.s. | |
| *Cyp7a1* | 12.776 | 0.728 | < 0.001 | 16.352 | 0.445 | < 0.001 | 3.576 |
| *Pparα* | 10.050 | 0.429 | < 0.001 | 12.177 | | n.s. | |
| *Srebf1* | 11.457 | | n.s. | 14.260 | | n.s. | |
| *Angptl8* | 19.961 | 0.445 | 0.044 | 1.538 | | n.s. | |
| *Pck1* | 9.856 | 2.546 | < 0.001 | 13.583 | | n.s. | |
| *G6pc* | 10.574 | | n.s. | 20.913 | | n.s. | |
| *Sirt1* | 13.856 | | n.s. | 14.276 | | n.s. | |
| *Sirt6* | 22.302 | | n.s. | 3.266 | | n.s. | |

1) n.s., not significant

**Supplementary Table 2** Primer sequences for qPCR

| *Gene* | Forward (5'→3') | Reverse (5'→3') | Size (bp) |
|---|---|---|---|
| *Per1* | agaagaaaacagcaccagct | tcttgagttataagaaccccaacatg | 98 |
| *Per2* | gccaagtttgtggagttcctg | cttgcaccttgaccaggtagg | 226 |
| *Bmal1* | gcagtgccactgactaccaaga | tcctggacattgcattgcat | 170 |
| *Rev-erbα* | ccctggactccaataacaacaca | gccattggagctgtcactgtag | 110 |
| *Sirt1* | cgggatatatttcctttgcaaactt | tatctatgctcgccttgcgg | 122 |
| *Sirt6* | gctggaggactgccacatta | atgtcggtgaattatgcagca | 138 |
| *Hsp70* | ccacaaaaccttaacatggaca | tgttccagtagcctgggaag | 165 |
| *p53* | tcttccagatgctcgggatac | gtcacagcacatgacggagg | 129 |
| *Wee1* | accccggagtttaacagagc | ctccgcacaagaccttcc | 110 |
| *c-Myc* | tccaagtaactcggtcatcatct | gctctccatccaatgttgcgg | 116 |
| *Angptl8* | ataagaatgcggccgcatggctgtgcttgctctctgcct | cgggatccggctgggagggctgctgtgtgga | 100 |
| *Srebf1* | atcggcgcggaagctgtcggggtagcgtc | actgtcttggttgttgatgagctggagcat | 116 |
| *G6pc* | tggtagccctgtctttctttg | ttccagcattcacactttcct | 90 |
| *Pck1* | gtgggcgatgacattgcc | actgaggtgccaggagcaac | 101 |
| *Pparα* | tgcaaacttggacttgaacg | aggaggacagcatcgtgaag | 80 |
| *Cyp7a1* | agcaactaaacaacctgccagtacta | gtccggatattcaaggatgca | 82 |
| *Hprt1* | atacaggccagactttgttggatt | tcactaatgacacaaacgtgattcaa | 110 |
| *Rplp0* | ctcactgagattcgggatatg | ctcccaccttgtctccagtc | 223 |

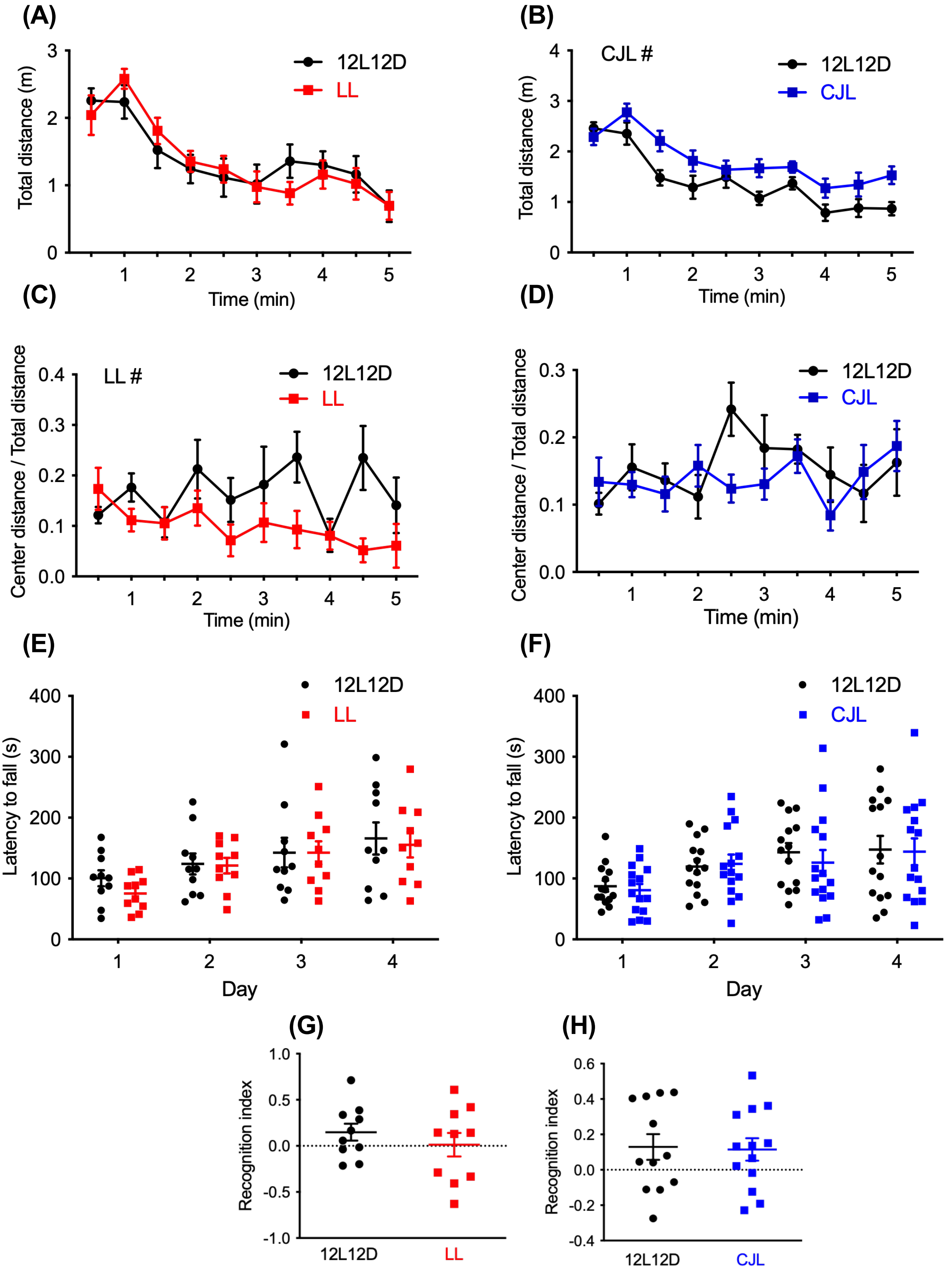


**Supplementary Figure 1** Constant light (LL) and chronic jet lag (CJL) had minimal effects on aging-associated behaviors. (A–D) Behaviors in the open field test. The total distance in the LL (A) or CJL (B) was evaluated as exploratory/spontaneous activity in a novel environment, while the center distance divided by the total distance in the LL (C) or CJL (D) was evaluated as anxiety-like behavior. (E, F) Latency to fall in the rotarod test in the LL (E) and CJL (F) groups as a measure of motor coordination. (G–J) Recognition index in the object recognition test in the LL (G) and CJL (H) as a measure of cognitive ability. Data are shown as mean ± standard error of the mean (S.E.M) (n = 10–15). #$p < 0.05$, repeated measures two-way ANOVA.

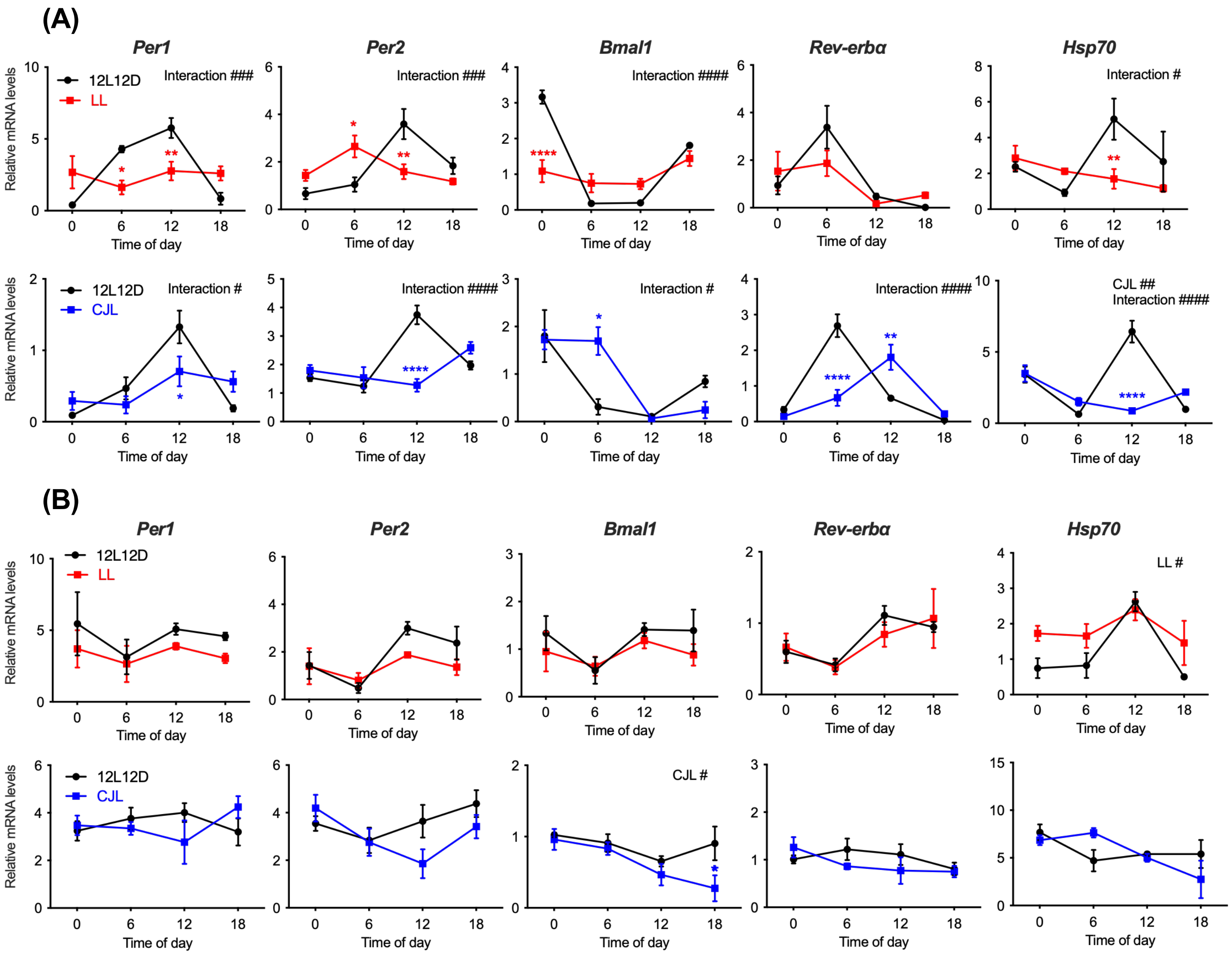


**Supplementary Figure 2** Constant light (LL) and chronic jet lag (CJL) differentially alter circadian gene expression in the liver and hippocampus. Expression of clock genes and *Hsp70* in the liver (A) and (B) hippocampus of mice under LL or CJL (n = 3–5). Mice in the control (12L12D) and CJL groups were transferred to constant darkness two days before euthanasia. The dark phase was initiated at 12:00 before implementing a constant darkness. Data are shown as mean ± standard error of the mean (S.E.M). #$p < 0.05$, ##$p < 0.01$, ###$p < 0.001$, ####$p < 0.001$, two-way ANOVA; *$p < 0.05$, **$p < 0.01$, ****$p < 0.0001$, Bonferroni's multiple comparison test.

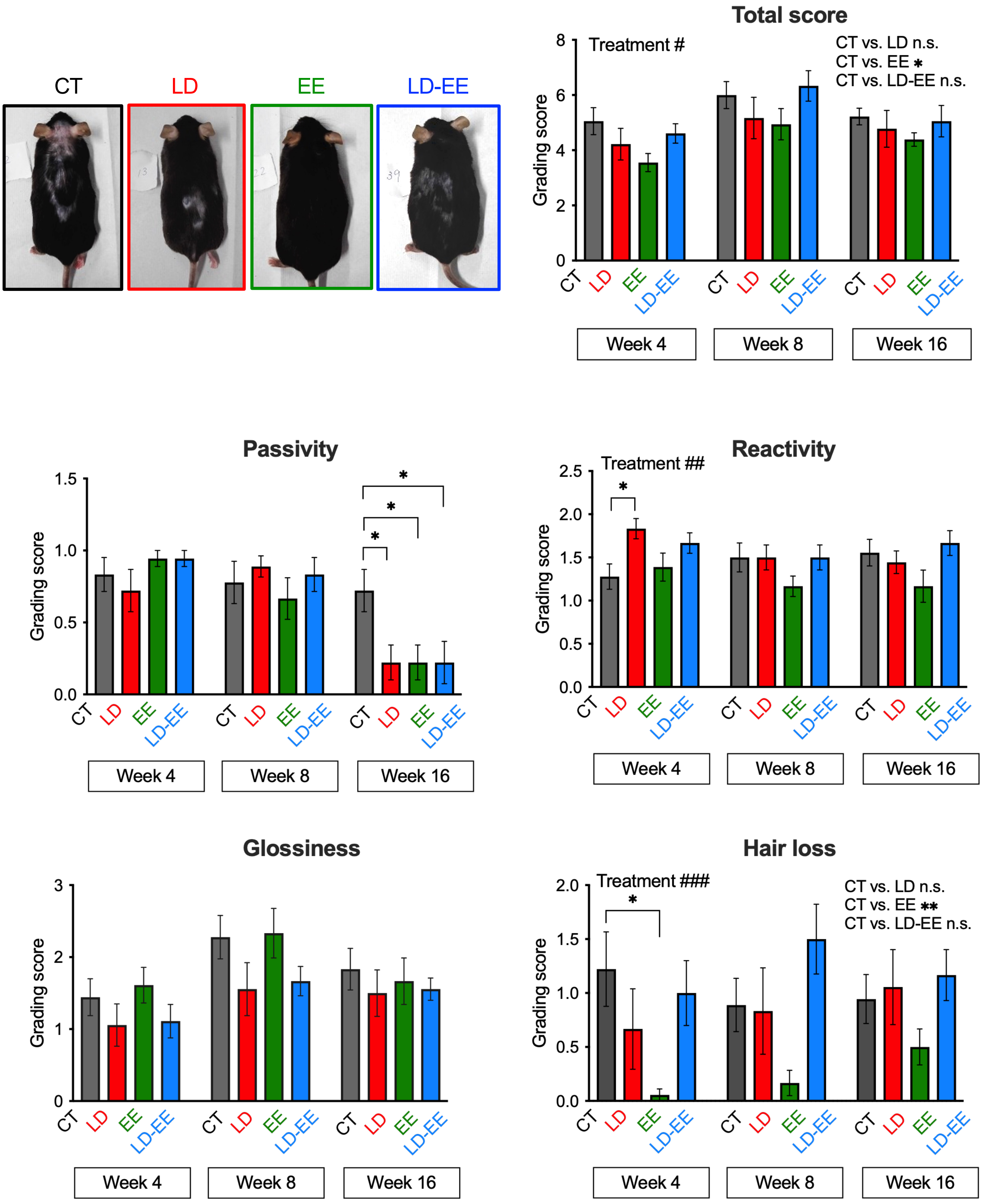


**Supplementary Figure 3** SJL-like conditions affected the grading scores. The grading scoring was performed according to a previous report[51]. Data are shown for control (CT), weekly shifts of light-dark cycles (LD), weekly housing in environmental enrichment (EE), and a combination of LD and EE (LD-EE) in weeks 4, 8, and 16, along with representative photos of mice in week 4. Data are shown as mean ± standard error of the mean (S.E.M) (n = 9). #$p < 0.05$, ##$p < 0.01$, ###$p < 0.001$, two-way ANOVA; *$p < 0.05$, Dunnett's multiple comparison test.

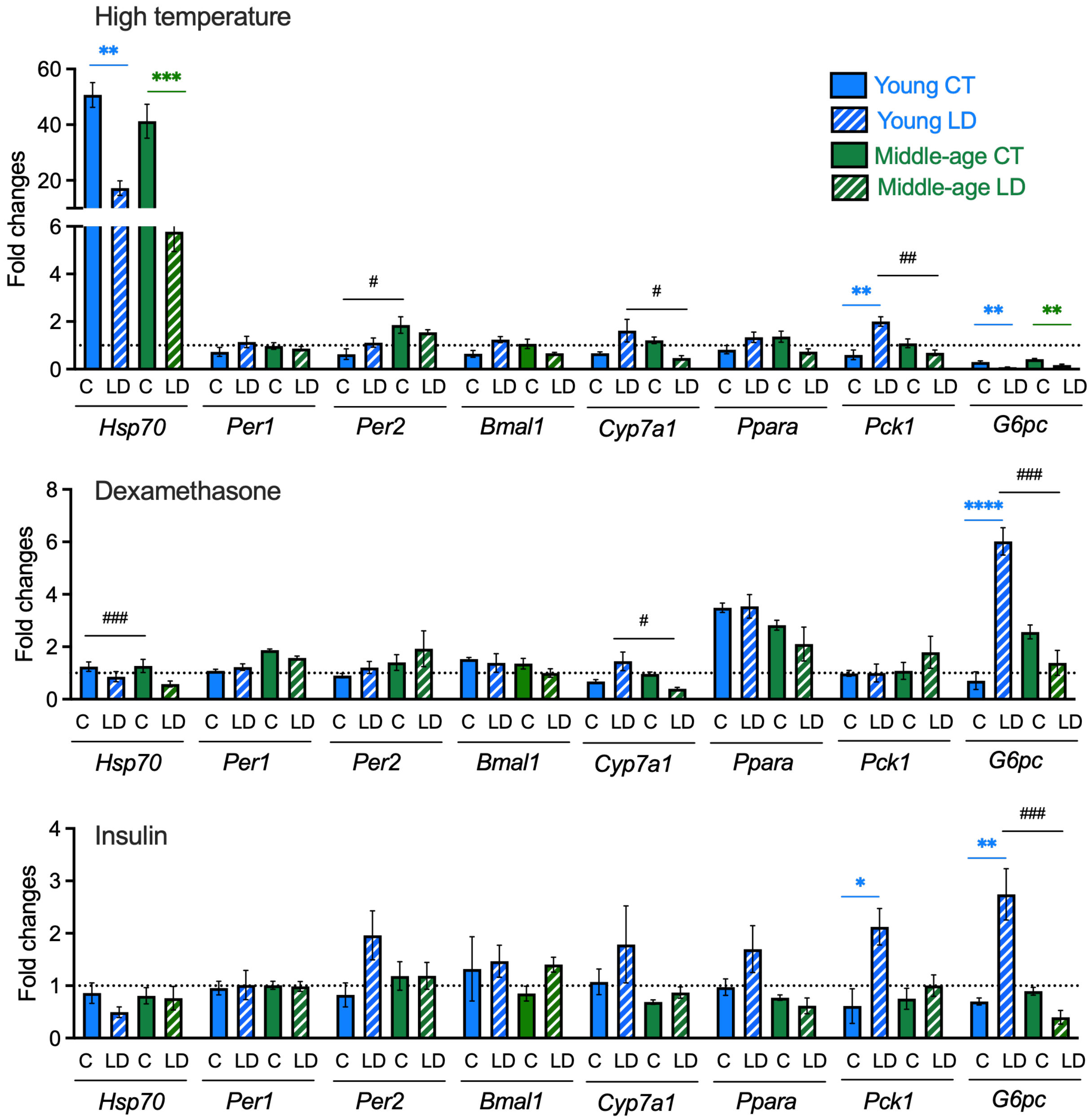


**Supplementary Figure 4** The responsiveness of gene expression in liver slices with 6 h of stimulation with high temperature, dexamethasone, or insulin. Liver slices were prepared from young and middle-aged mice kept under LD 12:12 (CT) or weekly shifts of light-dark cycles (LD). After 6h of stimulation, slices were analyzed for the expression of heat-sensitive *Hsp70*, circadian clock, and metabolic genes (three replicates per treatment). Data are shown as fold changes relative to the expression levels in non-stimulated slices. Data are shown as mean ± standard error of the mean (S.E.M). *$p < 0.05$, **$p < 0.01$, ***$p < 0.001$, ****$p < 0.0001$, Bonferroni's multiple comparison test, comparison between young CT and young LD and between middle-age CT and middle-age LD. #$p < 0.05$, ##$p < 0.01$, ###$p < 0.001$, Bonferroni's multiple comparison test, comparison between young and middle-aged CT groups and between young and middle-aged LD groups.
